# Reassessment and typification of *Opuntia canterae* (Opuntioideae, Cactaceae), an endemic prickly-pear cactus of Uruguay

**DOI:** 10.1101/2020.03.06.981522

**Authors:** Matias Köhler, Lucas C. Majure

## Abstract

*Opuntia* is the most widespread genus of Cactaceae, naturally occurring throughout arid and semi-arid areas of the Americas. Many of the species have taxonomic problems owed to incomplete original descriptions, lack of type designations, a paucity of taxonomic revisions and in general, difficult species delimitation resulting from hybridization, morphological plasticity and poor specimen preparation. However, efforts are being undertaken to fill in the gaps in our distributional, morphological and phylogenetic knowledge of the group. Here, we reassess the name *Opuntia canterae*, providing an updated description, typification, photographs, distribution map, conservation assessment and additional notes.

**Material and methods:** Extensive fieldwork was conducted, along with comprehensive herbarium and literature review. Morphological characters were assessed based on the commonly used characters used for prickly pears. Species delimitation is proposed based on our morphological studies, taxonomic and literature revision, as well as preliminary phylogenetic studies. The IUCN guidelines were followed to provide a conservation assessment of the species.

**Key results:** *Opuntia canterae* is reassessed as a distinct species separated from its previous synonym (*O. elata*) by the elliptic to long-oblanceolate stem segments, acute to conical flower bud apex and long-obconic fruits. An epitype is here designate for purposes of the precise designation of the name to the taxon. The species is considered endemic to Uruguay and is assessed as Vulnerable (VU) using IUCN criteria, but more fieldwork will be necessary to provide a more precise conservation status.

## Introduction

*Opuntia* Mill. is the most widespread genus of Cactaceae, naturally occurring from southern South America (Argentina) to northern North America (Canada) (Britton and Rose 1919; Anderson 2001; Majure et al. 2012). The group has a putative origin during the Late Miocene (∼7- 5 Mya) in southwestern South America with subsequent dispersal events of lineages to northern South America, the Caribbean region, Central America and to the North American deserts (Arakaki et al. 2011; Majure et al., 2012). The group exhibits a variety of morphological characters such as a shrubby or tree growth form, dry/fleshy fruit, epidermis and seeds either pubescent or glabrous, dioecy/monoecy, ornithophilic/melittophilic pollination, as well as other discrete and phenotypically plastic characters (Schumann 1899, Britton and Rose 1919, Backeberg 1958, Anderson 2001, Hunt et al. 2006, Majure and Puente 2014, Majure et al. 2017).

Eight major clades have been recovered within *Opuntia* s.str. (Majure et al, 2012; Köhler et al., *in prep.*), and the South American species are mainly nested in two of these clades: *Macbridei* and *Elatae* (sensu Majure et al. 2012, Köhler et al. *in prep.*). The *Macbridei* clade includes species occurring in the northern part of South America, from central Peru to central Colombia (Britton and Rose 1919, Anderson 2001, Madsen 1989, Vega 2013, Majure and Puente 2014), while the *Elatae* clade includes the southern South American lineages occupying mainly the Pampa and the Chaco regions, as well as the Galapagos Island species (Britton and Rose 1919, Leuenberger 2002, Majure et al. 2012, Font 2014, Las Peñas et al. 2017, Köhler et al. 2018, Köhler et al. *in prep.*).

Some of the southern South American (sSA) species of *Opuntia* have a confused taxonomic history. Many of these taxa were described based on materials collected by Old World naturalists that were travelling to the New World and sending biological materials to European gardens (Pontes et al. 2017). That routine led to many names, which were poorly described, based on morphotypes grown under greenhouses conditions, with insufficient diagnoses or use of characters and usually without the designation of nomenclatural types (Haworth 1812; Pfeiffer 1837; Salm-Dyck 1850). Beyond that, many European naturalists that migrated to the New World and started to contribute to the knowledge of local floras also often failed to cite original materials or provide precise descriptions of the novel species proposed (Spegazzini 1899, 1901, 1902, 1905, 1925; Arechavaleta 1905). Just recently, enormous efforts have been made to better assess the identity and the interpretation of many of these names with typifications and a handful of taxonomic revisions (Crook & Mottram 1995, 1996, 1997, 1998, 1999, 2000, 2001, 2002, 2003, 2004, 2005; Leuenberger 1993, 2001a, 2001b, 2002; Font 2014; Las Peñas et al. 2017).

*Opuntia canterae* was described by Arechavaleta (1905) as a distinct species based on his knowledge of the Uruguayan flora and neighbouring areas. The description included a comprehensive diagnosis with a complementary description accompanied by personal observations about the ecology and distribution of the species (Arechavaleta 1905). This taxon was further treated as a doubtful species by Britton & Rose (1919), which merely transcribed the original description of Arechavaleta without mentioning the detailed knowledge about the ecology, etc., of the species. Bertram (1929, 1931) reported his success in growing *Opuntia* species in Germany, illustrating a flowering prickly pear identified as *O. canterae* by Hern W. Weingart. Herter (1957) included the species in his study of the Uruguayan flora and illustrated the species with narrow and spineless stem segments, with visibly pointed apex flower buds. Backeberg (1958) reproduced Arechavaleta’s description providing a photograph of ambiguous assignment, without providing any additional information. Anderson (2001) listed the species in his treatment based on the previous, sparse descriptions. Leuenberger (2002), in the first attempted taxonomic revision of a series from the sSA species of *Opuntia* (series *Armatae* K. Schum. = *Elatae* Britton & Rose), was unable to critically access the identity of the taxon, and suggested that it may belong in *O. elata* Salm-Dyck or *O. cardiosperma* K.Schum.

Font (2014), in an attentive revision of the series *Armatae*, proposed a novel set of morphological characters for a comprehensive circumscription of the species. Besides the already used morphology of the stem segments (cladodes), spination and habit of the species, Font (2014) introduced the morphology of the flower bud apices, stigma colour and the inner pericarpel tissue colour as useful characters to diagnose taxonomic entities that have been problematic historically. Even so, Font (2014) suggested *O. canterae* as synonym of a broadly circumscribed *O. elata*, and later Las Peñas et al. (2017) retained it in the synonymy of *O. elata.*

During a broad taxonomic, systematic and floristic revision of the southern South American species of *Opuntia*, a distinct morphotype have been observed in the Pampean region of Uruguay, and further analyses were carried out to assess the identity for those materials, which conform to *Opuntia canterae*. Here, we propose a reassessment of *O. canterae*, providing a typification, an updated description, photographs, conservation assessment and additional notes about the species.

## Material and Methods

Extensive field trips were carried out in southern South America encompassing the main natural ecoregions to obtain data about natural populations of *Opuntia*. The region is represented by subtropical grasslands permeated by rocky outcrops that compose the Pampa or Río de La Plata grassland (Andrade et al., 2018) and the Chaco region (Pennington et al., 2000). Major herbaria from the region have been revised to check distribution records and specimen identification of all *Opuntia*: BA, BAF, BCWL, CORD, CTES, HAS, ICN, LIL, LP, MCN, MVFA, MVJB, MVM, SI (herbarium abbreviations following Thiers (2020+, continuously updated), except BCWL, non-indexed herbarium of the Biological Control of Weeds Laboratory (FuEDEI), Hurlingham, Buenos Aires, Argentina). The digital database of Brazilian collections was also consulted through the SpeciesLink platform (2019) to check herbaria from disparate geographical regions.

A literature review was carried out comprising the main magnum opus that contains description of southern South American *Opuntia* species (Arechavaleta 1905; Spegazzini 1901, 1905, 1925; Schumann 1890, 1899a,b; Britton and Rose 1919; Backeberg 1958, 1966; Ritter 1979, 1980), as well as recent revisions, floras and taxonomic treatments (Kiesling 1999, 2005; Kiesling and Ferrari 2005; Kiesling et al. 2008; Machado et al. 2008; Leuenberger 2002; Font 2014; Las Peñas et al. 2017). The morphological characters used for identification of the southern South American species of *Opuntia* followed those proposed by Font (2014) and Las Peñas et al. (2017), which have been reported as useful for species delimitation in other sSA *Opuntia* species (Köhler et al. *under review*). For the conservation status assessment of the species, the GeoCAT tool (Bachman et al. 2011) was used to evaluate the area of occupancy (AOO) and the extent of occurrence (EOO), using a cell width of 5 km based on our observations. The criteria followed those proposed by the IUCN Red List (IUCN, 2019).

## Results and Discussion

*Opuntia canterae* has been treated as a doubtful taxon (Britton & Rose 1919; Leuenberger 2002; Kiesling et al. 2008), valid species (Anderson 2001), or more recently as synonym of *O. elata* (Font 2014; Las Peñas et al. 2017). During our recent field expeditions, a distinct morphotype has been observed in the Pampean region of Uruguay, and none of the previous taxonomic treatments included its morphological features under the circumscription of the species proposed, nor within the identification key provided. The combined features in *O. canterae* of the elliptic to long-oblanceolate stem segments, acute flower bud apices and long-obconic ripe fruits (Fig. 1), separate the species from *Opuntia elata*, which includes specimens with obovate-oblong stem segments, rounded/globose flower bud apices and pyriform fruits. Our preliminary phylogenetic studies of the sSA species of *Opuntia* (Köhler et al., *in prep.*) reinforces *O. canterae* as a distinct evolutionary lineage of the *Elatae* clade (sensu Majure et al. 2012), which led us to propose a reassessment of the species.

**Figure 1.**
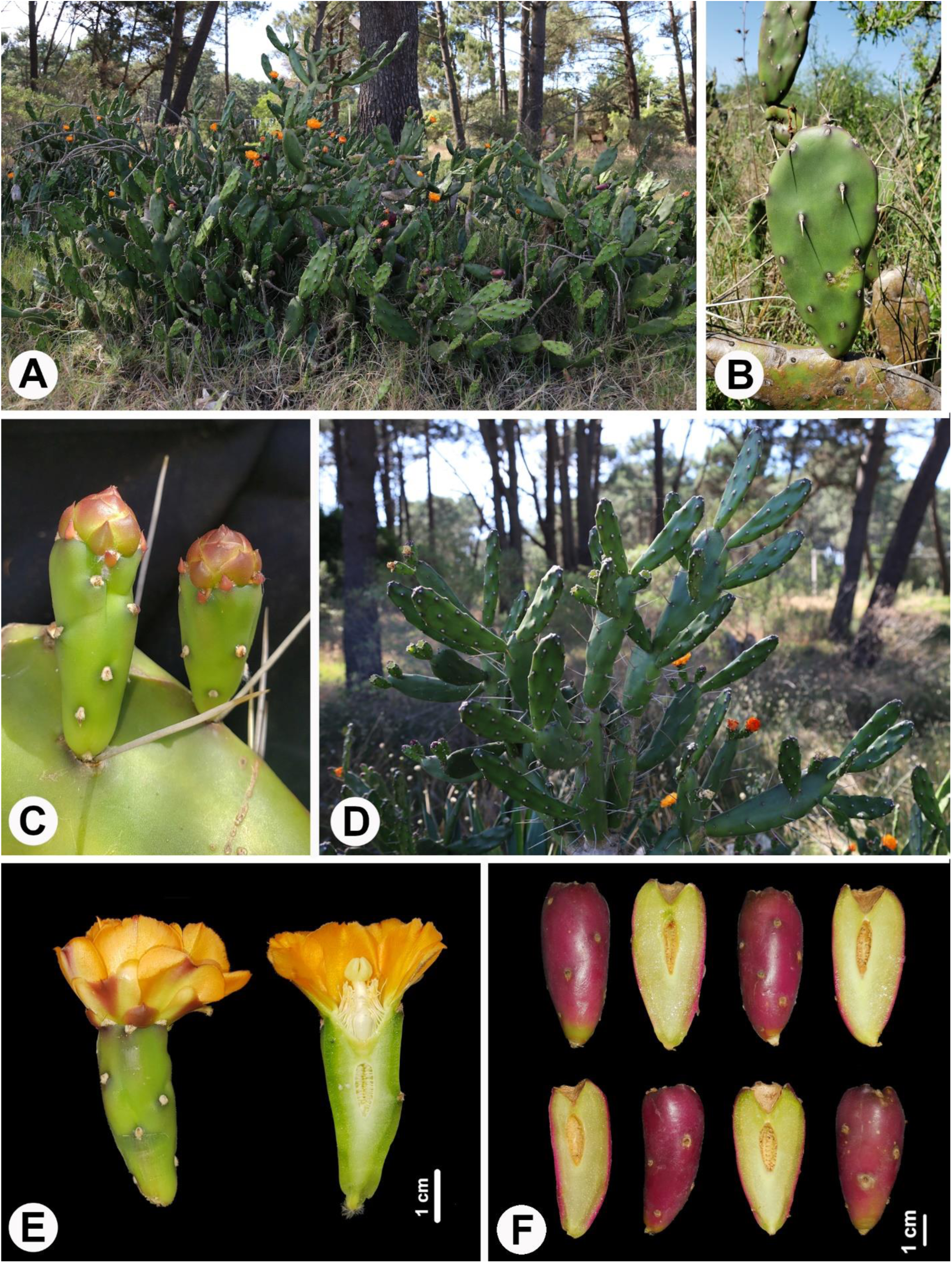
Morphological features of *Opuntia canterae.* **A**. Plant in habitat (*M. Köhler 316*). **B.** Detailed stem segment resembling morphotype designated as neotype (*M. Köhler 550*). **C.** Detail of the acute flower bud apex (*M. Köhler 550*). **D.** Elliptic to long-oblanceolate stem segments, showing growing cladodes with protuberant areoles encircled with dark-violet coloration from betalain pigmentation (*M. Köhler 316*). **E.** Flower in transverse section showing orange tepals, obconic pericarpel, sterile stamens and obovate to elliptical ovary (*M. Köhler 550*). **F.** Transverse section of the long-obconic dark-purple ripe fruits showing the sterile ovaries and light green inner pericarpel tissue (*M. Köhler 316*).

### Taxonomic treatment

***Opuntia canterae*** Arechav., *Anales del Museo Nacional de Montevideo, Tomo II* (Arechavaleta 1905: 278–280, as *O. canterai*) Figs. 1–4.

**Type**: Not designated; **Neotype:** designated by Las Peñas et al. 2017, Lám. LX in Osten (1941); **Epitype** (**designated here**): URUGUAY. Canelones: Neptunia, 34°47’2.73”S, 55°53’11.75”W, 6 December 2017, *M. Köhler et al. 316* (ICN 201773, barcode 00043878, isoepitype: MVM).

<u>Shrub</u>, erect, 1.5–2(>2) m tall. <u>Stem segments (cladodes)</u> 13–30 x 4–6 cm, 2–2.5(–3) cm thick, elliptic to long-oblanceolate, dark green, apex rounded to obtuse, base attenuate, occasionally forming subterete proximal stems. <u>Areoles</u> 6–9 at midsection of cladode, 0.4–0.6 cm in diameter, circular to elliptic, frequently protuberant on growing cladodes, circled with dark-violet stains. <u>Leaves</u> conic, dark-violet, 3–4 mm, usually only on the apex of new cladodes or pericarpel, quickly ephemeral. <u>Glochids</u> present but not developed, poorly exerted outside of the areoles, ferruginous. <u>Spines</u> 0–1(–2) per areole, acicular, white to light grey, reflexed (when < 3 cm) to straight (when > 4–10 cm). <u>Pericarpel</u> 3.5–4 x 1.5(–2) cm, obconic. <u>Flower bud apex</u> acute to conical, external tepals reddish, obcordate with mucronulate apex; inner tepals orange, largely obovate with mucronulate apex; flower at anthesis 3–5 cm in diameter. <u>Stamens</u> numerous with pale yellow filaments and anthers when present; frequently sterile. <u>Stigma</u> 6–7 lobed, connivent, cream-colored. <u>Style</u> cylindric to obclaviform, 1.7–2 x 0.3–0.5 cm. <u>Ovary</u> 1–1.3 x 0.4–0.7 cm, obovoid, in the upper third of the pericarpel. <u>Fruit</u> 5.5-7 x 2.5-3 cm, long-obconic, red to dark-purple when ripe, inner pericarpel light greenish. <u>Seeds</u> flattened (not seen in recent specimens).

#### Specimens examined — URUGUAY

Montevideo, Pocitos, December 1921, *C. Osten 16016* (MVM). San José: Rincón del Pino, 34°30’8.61”S, 56°50’7.37”W, 4 December 2017, *M. Köhler et al. 299* (ICN), *M. Köhler et al. 302* (ICN); Libertad, 34°39’48.17”S, 56°35’3.69”W, 4 December 2017, *M. Köhler et al. 303* (ICN). Canelones: Neptunia, 34°47’2.73”S, 55°53’11.75”W, 6 December 2017, *M. Köhler et al. 316* (ICN). Río Negro: Nuevo Berlin, 32°53’10.9”S, 58°02’42.4”W, 23 January 2020, *M. Köhler et al. 550* (ICN).

#### Distribution

Only recorded in Uruguay near Río de la Plata and Río Uruguay (Esteros and Algarrobales del Río Uruguay) in the departments of Canelones, Río Negro, San José and Montevideo (Fig. 2).

**Figure 2.**
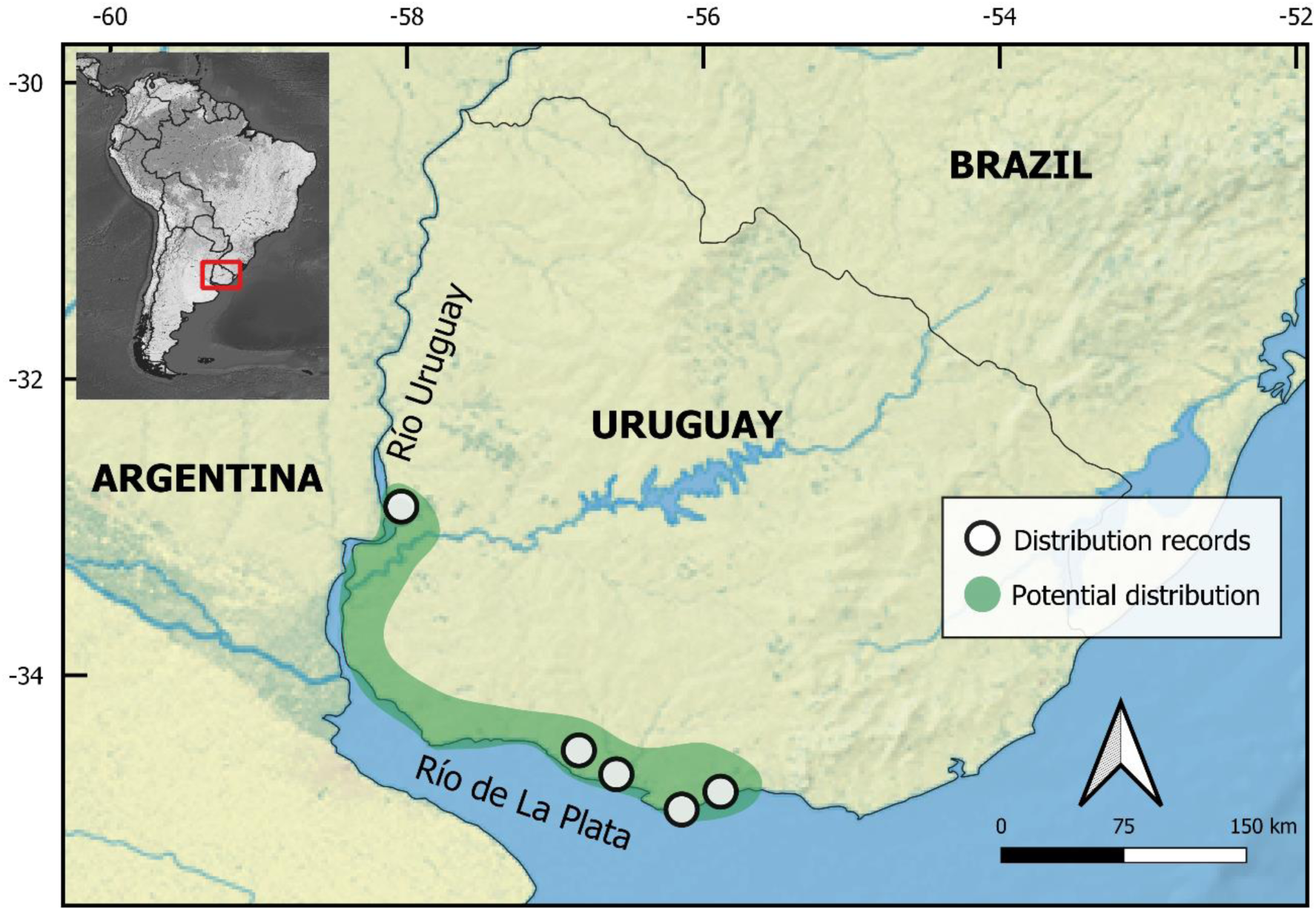
Distribution map of *Opuntia canterae.* The white dots indicate the known records of distribution, while the green area indicate a potential distribution of the taxon that must be further investigated.

#### Habitat

The species is originally described as occurring along the Uruguayan coastal plain of the “Río de La Plata” river, on sandy or rocky (granite) soils, where it has been sparsely observed. New records have been observed in the extreme northwest part of the “Río de La Plata”, on the margins of the “Río Uruguay”, in the “Esteros y Algarrobales del Río Uruguay”, suggesting a broader distribution that must be further investigated.

#### Conservation assessment

The currently known records of the species are reduced to only four localities. Although many of the natural areas of Uruguay has been converted to agroindustry plantations of *Eucalyptus* spp., *Glycine max* (soybean) or to smallholder livestock ranching and annual agriculture, we expect that there are more localities where the species occurs but have not been reported yet. Based on the known distribution, the extent occurrence (EOO) of the species is estimated to be ∼6,400 km^2^, which places it under the Vulnerable (VU) category, whereas its area of occupancy (AOO) is estimated to be 100 km^2^, which put it under the category of Endangered (EN) using the subcriterion B2a of IUCN (2019). Admitting that there are still lacking appropriate data to make an adequate assessment of its risk of extinction, a Data Deficient category (DD) would be most appropriate for this taxon. However, considering that the species has long been ignored as a doubtful taxa or synonym of *O. elata/O. cardiosperma*, with few known records, and lives in an highly threatened environment, we are giving a precautionary IUCN Red List Category of Vulnerable: VU B1a,b(ii,iii)+2a,b(iii), suggesting that more fieldwork is necessary to critically evaluate the conservation status of this species.

#### Phylogenetic relationships

This species was not sampled in previous phylogenetic analyses (Majure et al. 2012, Majure and Puente 2014, Realini et al. 2014, Majure et al. 2020). However, newly generated data has revealed the species as a distinct lineage in the *Elatae* clade (Köhler et al., *in prep.*, sensu Majure et al. 2012), being closely related with some species treated in series *Armatae* K.Schum. such as *O. elata* and *O. megapotamica* (sensu Font 2014).

#### Notes

Las Peñas et al. (2017) designated a neotype based on a photographic plate provided by Osten (1941, LAM. LX). The same photography was found in the MVM herbarium on a duplicate herbarium sheet, with one them being accompanied by personal notes of C. Osten (Fig. 3) which were almost entirely transcribed in Osten (1941). Our field studies allowed us to observe the same features provided by the photograph, as well as the original descriptions of Arechavaleta (1905), in those populations sampled (Fig 1A–B, D). However, considering that the neotype is a photograph of a putative juvenile plant, lacking important characters to be critically identified, we designate here an epitype containing features applied for the precise designation of the name of the species (Fig. 4). The species is still lacking knowledge about its biology. As pointed in Arechavaleta (1905) and confirmed in our field work, *O. canterae* frequently has sterile stamens and fruits, thus, it will be necessary to further investigate putative dioecy in this species, as reported for other *Opuntia* species (Díaz & Cocucci 2008; Majure & Puente 2014). Additionally, chromosome counts must be explored in *O. canterae*, since there are no information about it ploidy level yet.

**Figure 3.**
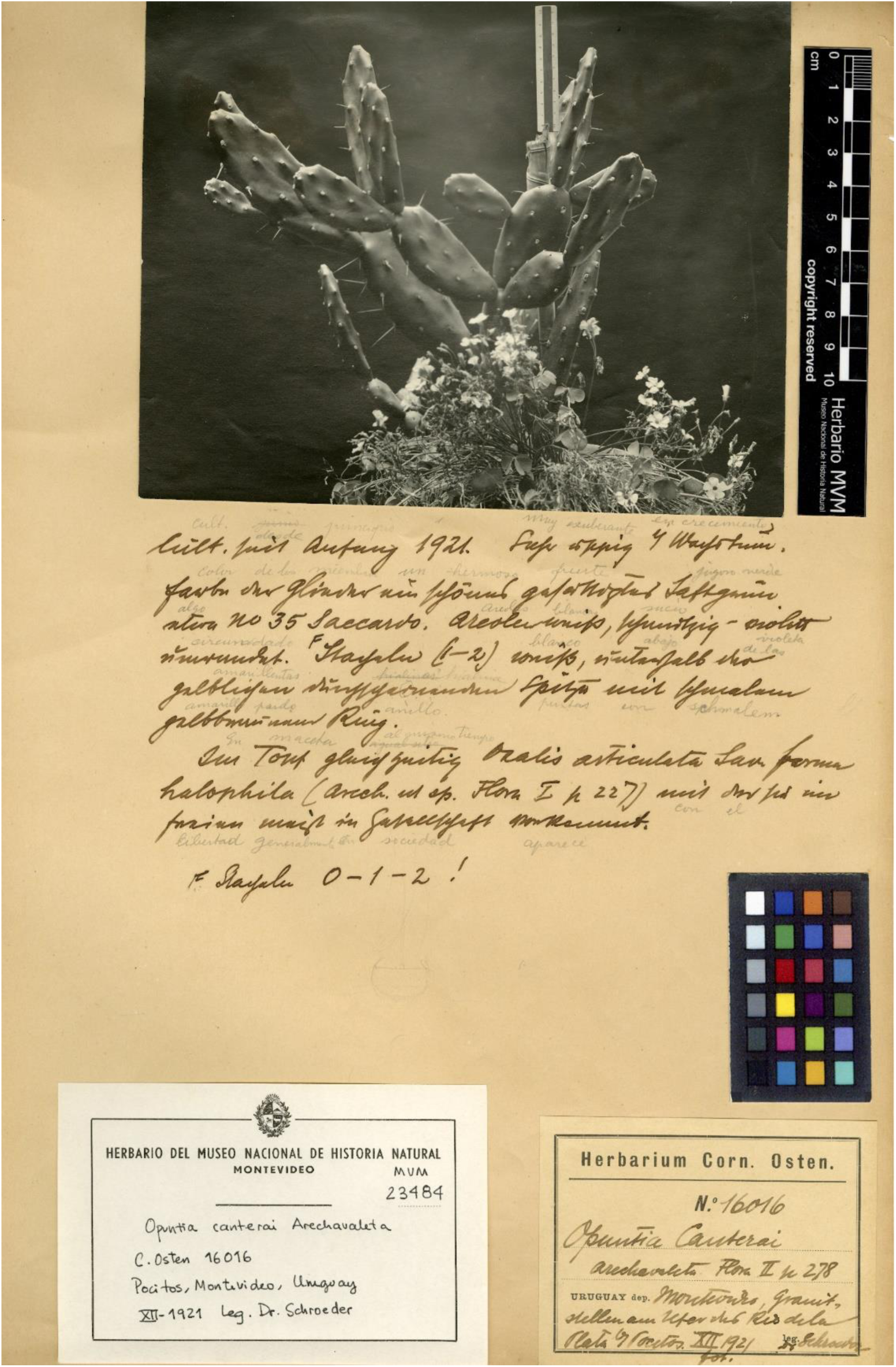
Herbarium specimen from Cornelius Osten Herbarium (MVM 23484, *C. Osten 16016*), which includes the photograph designated as the neotype by Las Peñas et al. (2017), accompanied by personal notes from C. Osten.

**Figure 4.**
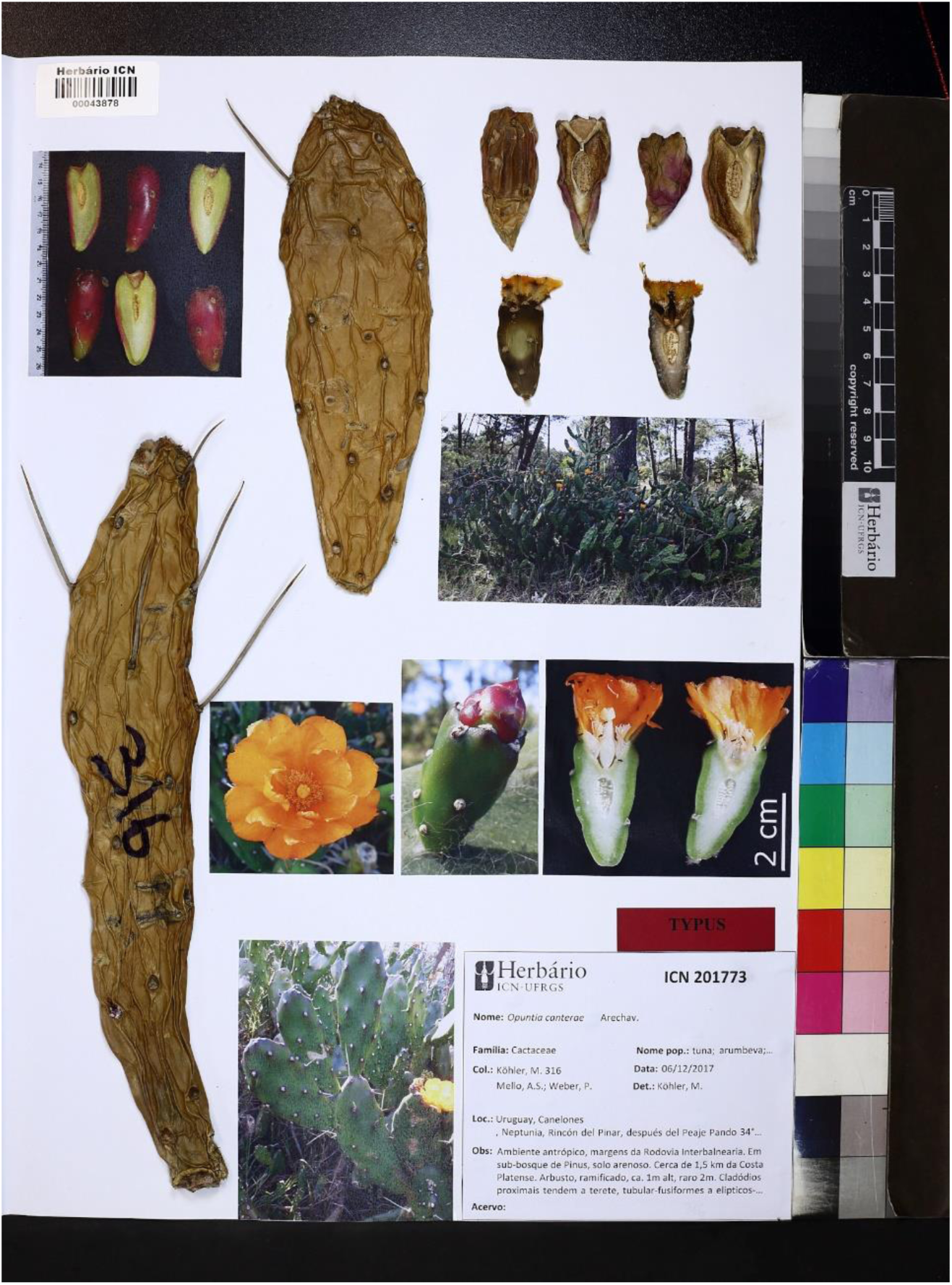
Epitype of *Opuntia canterae* (ICN 201773, barcode 00043878, *M. Köhler et al. 316*), which includes important characters to critically apply the name to the taxon, such as the elliptic to long-oblanceolate stem segments, acute flower bud apices and long-obconic fruits.

## Acknowledgements

We thanks to Dr. Jefferson Prado (Instituto de Botânica, Herbarium SP) for providing helpful comments about nomenclature and typification; Anderson S. Mello, Philipy Weber, Lucas Kaminski, Luis Volkmann and Curtis Callaghan for providing help during fieldwork; Ivan Grela (UPM) for permission of studies in the protected area of the “Esteros y Algarrobales del Río Uruguay (Río Negro)” in Uruguay; Andrés González and Meica Valdivia for helping digitalizing MVM material and permission for use it; and the staff from ICN herbarium for support in digitalizing the epitype. MK is grateful to the American Society of Plant Taxonomists (ASPT), Cactus and Succulent Society of America (CSSA), International Association for Plant Taxonomy (IAPT) and IDEA WILD for supporting part of the research here reported; MK also thanks the Brazilian National Council for Scientific and Technological Development (Conselho Nacional de Desenvolvimento Científico e Tecnológico - CNPq) for his PhD scholarship, and the PDSE/CAPES for support his period as Visiting Researcher at the Florida Museum of Natural History (FLMNH, University of Florida, UF, USA). This study was also financed in part by the Coordenação de Aperfeiçoamento de Pessoal de Nível Superior – Brasil (CAPES) - Finance Code 001 and start-up funds to LCM from UF and the Florida Museum of Natural History.

## References

Anderson E.F. (2001) The cactus family. Portland, Timber Press.

Andrade B.O., Marchesi E., Burkart S., Setubal R.B., Lezama F., Perelman S., Schneider A.A., Trevisan R., Overbeck G.E., Boldrini I.I. (2018) Vascular plant species richness and distribution in the Río de la Plata grasslands. Botanical Journal of the Linnean Society 188: 250–256. https://doi.org/10.1093/botlinnean/boy063

Arakaki M., Christin P.-A., Nyffeler R., Lendel A., Eggli U., Ogburn R.M., Spriggs E., Moore M.J., Edwards E.J. (2011) Contemporaneous and recent radiations of the world’s major succulent plant lineages. Proceedings of the National Academy of Sciences 108: 8379–8384. https://doi.org/10.1073/pnas.1100628108

Arechavaleta J. (1905) Anales del Museo Nacional de Montevideo. Volumen 5, Tomo II. Montevideo.

Bachman S., Moat J., Hill A., Torre J. de la, Scott B. (2011) Supporting Red List threat assessments with GeoCAT: geospatial conservation assessment tool. ZooKeys 150: 117–126. https://doi.org/10.3897/zookeys.150.2109

Backeberg C. (1958) Die Cactaceae, Band I. Jena, Gustav Fischer Verlag.

Backeberg C. (1966) Das Kakteenlexikon. Stuttgart, Gustav Fischer Verlag.

Bertram P. (1929) *Opuntia canterai* Arech. Monatsschrift der Deutschen Kakteen-Gesellschaft 1(12): 239–241.

Bertram P. (1931) *Opuntia canterai* Arech. Monatsschrift der Deutschen Kakteen-Gesellschaft 3(11):259–260

Britton N.L., Rose J.N. (1919) The Cactaceae. Washington, Carnegie Institute.

Crook R., Mottram R. (1995) *Opuntia* Index Part 1: Introduction and A-B. Bradleya 1995: 88–118. https://doi.org/10.25223/brad.n13.1995.a10

Crook R., Mottram R. (1996) *Opuntia* Index Part 2: Nomenclatural note and C-E. Bradleya 1996: 99–144. https://doi.org/10.25223/brad.n14.1996.a15

Crook R., Mottram R. (1997) *Opuntia* Index Part 3: Nomenclatural note and F. Bradleya 1997: 98–112. https://doi.org/10.25223/brad.n15.1997.a12

Crook R., Mottram R. (1998) *Opuntia* Index Part 4: G-H. Bradleya 1998: 119–136. https://doi.org/10.25223/brad.n16.1998.a11

Crook R., Mottram R. (1999) *Opuntia* Index Part 5: Nomenclatural note and I-L. Bradleya 1999: 109–131. https://doi.org/10.25223/brad.n17.1999.a8

Crook R., Mottram R. (2000) *Opuntia* Index Part 6: M-O. Bradleya 2000: 113–140. https://doi.org/10.25223/brad.n18.2000.a9

Crook R., Mottram R. (2001) *Opuntia* IndexPart 7: Nomenclatural note and P–Q. Bradleya 2001: 91–116. https://doi.org/10.25223/brad.n19.2001.a11

Crook R., Mottram R. (2002) *Opuntia* IndexPart 8: R. Bradleya 2002: 51–66. https://doi.org/10.25223/brad.n20.2002.a9

Crook R., Mottram R. (2003) *Opuntia* Index Part 9: S. Bradleya 2003: 63–86. https://doi.org/10.25223/brad.n21.2003.a13

Crook R., Mottram R. (2004) *Opuntia* Index Part 10: T-V. Bradleya 22: 53–76. https://doi.org/10.25223/brad.n22.2004.a7

Crook R., Mottram R. (2005) *Opuntia* Index Part 11: W-Z, cultivars. Bradleya 2005: 57–78. https://doi.org/10.25223/brad.n23.2005.a8

Díaz L., Cocucci A.A. (2008) Functional gynodioecy in *Opuntia quimilo* (Cactaceae), a tree cactus pollinated by bees and hummingbirds. Plant Biology 5(5): 531–539. https://doi.org/10.1055/s-2003-44783

Font F. (2014) A revision of *Opuntia* series Armatae K. Schum. (Opuntia ser. Elatae Britton & Rose) (Cactaceae - Opuntioideae). Succulent Plant Research 8: 51–94.

Haworth A. H. (1812) Synopsis plantarum succulentarum. London, Richardi Taylor et Soch. https://doi.org/10.5962/bhl.title.9462

Herter G.G. (1957) Estudios botánicos en la región Uruguaya. 14(2): without pagination. Montevideo.

Hunt D., Taylor N., Charles G. (2006) The New Cactus Lexicon. Milborne Port, DH Books.

IUCN (2019) Guidelines for using the IUCN Red List Categories and Criteria. Version 13. Prepared by the Standards and Petitions Sub-Committee. Available at https://cmsdocs.s3.amazonaws.com/RedListGuidelines.pdf [accessed 15 Dec. 2019].

Kiesling R. (1999) Cactaceae. In: Zuloaga F.O., Morrone, O. (eds.) Catálogo de las plantas vasculares de la República Argentina II: Dycotyledoneae. Monographs in Systematic Botany 74: 423–489. St. Louis, Missouri Botanical Garden Press.

Kiesling R. (2005) Cactaceae. In: Trancoso N.S., Bacigalupo, N.M. (eds.) Flora ilustrada de Entre Ríos (Agentina). Dicotiledoneas Arquiclamídeas. B. Geraniales a Umbelifloriales. Buenos Aires, INTA.

Kiesling R., Ferrari O. (2005) 100 cactus Argentinos. Buenos Aires, Albatros.

Kiesling R., Larocca J., Faúndez L., Metzing D., Albesiano S. (2008) Cactaceae: *Opuntia.* In: Zuloaga F., Morrone O., Belgrano M. (eds.) Catálogo de las plantas vasculares del cono sur. Vol. 2. Dycotyledoneae: Acanthaceae-Fabaceae. Monographs in Systematic Botany 107. St. Louis, Missouri Botanical Garden Press.

Köhler M., Font F., Souza-Chies T.T. (2018) First record of *Opuntia rioplatense* (Cactaceae) for the Brazilian Flora. Phytotaxa 379(4): 293–296. http://dx.doi.org/10.11646/phytotaxa.379.4.3

Las Peñas M.L., Oakley L., Moreno N.C., Bernardello G. (2017) Taxonomic and cytogenetic studies in *Opuntia* ser. Armatae (Cactaceae). Botany 95: 101–120. https://doi.org/10.1139/cjb-2016-0048

Leuenberger B.E. (1993) Interpretation and typification of *Cactus opuntia* L., *Opuntia vulgaris* Mill., and *O. humifusa* (Rafin.) Rafin. (Cactaceae). Taxon 42: 419–429. https://doi.org/10.2307/1223152

Leuenberger B.E. (2001a) *Opuntia paraguayensis* (Cactaceae) reassessed. Willdenowia 31: 181–187. https://doi.org/10.3372/wi.31.31116

Leuenberger B.E. (2001b) The type specimen of *Opuntia cardiosperma* (Cactaceae), new synonyms and new records from Argentina and Paraguay. Willdenowia 31: 171–179. https://doi.org/10.3372/wi.31.31115

Leuenberger B.E. (2002) The South American Opuntia ser. Armatae (= O. ser. Elatae) (Cactaceae). Botanische Jahrbücher fur Systematik, Pflanzengeschichte und Pflanzengeographie 123 (4): 413–439.

Machado M. (2008) Notes on Brazilian taxa of series Armatae (Elatae). Cactaceae Systematics Initiatives 24: 33–35.

Madsen J.E. (1989) Opuntia. In: Harling G., Andersson L. (eds) Flora of Ecuador 45. Cactaceae. Berlings, Azlov.

Majure L.C., Judd W.S., Soltis P.S., Soltis D.E. (2017) Taxonomic revision of the *Opuntia humifusa* complex (Opuntieae: Cactaceae) of the eastern United States. Phytotaxa 290: 1–65. https://doi.org/10.11646/phytotaxa.290.1.1

Majure L.C., Puente R. (2014) Phylogenetic relationships and morphological evolution in *Opuntia* s. str. and closely related members of tribe Opuntieae. Succulent Plant Research 8: 9–30.

Majure L.C., Puente R., Griffith M.P., Judd W.S., Soltis P.S., Soltis D.E. (2012) Phylogeny of *Opuntia* s.s. (Cactaceae): clade delineation, geographic origins, and reticulate evolution. American Journal of Botany 99: 847–864. https://doi.org/10.3732/ajb.1100375

Majure L.C., Köhler M., Font F. (2020) North American Opuntias (Cactaceae) in Argentina? Remarks on the phylogenetic position of *Opuntia penicilligera* and a closer look at *O. ventanensis*. Phytotaxa 428 (3): 279–289. http://dx.doi.org/10.11646/phytotaxa.428.3.9

Pennington R.T., Prado D.E., Pendry C.A. (2000) Neotropical seasonally dry forests and Quaternary vegetation changes. Journal of Biogeography 27: 261–273. https://doi.org/10.1046/j.1365-2699.2000.00397.x

Pfeiffer L.G.K. (1837) Enumeratio diagnostica Cactearum hucusque cognitarum. Berlin, L. Oehmigke. https://doi.org/10.5962/bhl.title.15207

Pontes R.C., Marchiori J.N.C., Neto L.W. (2017) Notas históricas sobre a família Cactaceae no Rio Grande do Sul (Brasil) e Uruguai. I – Período Clássico (1818-1950): viajantes naturalistas e botânicos europeus. Balduinia 0: 01–11. https://doi.org/10.5902/2358198026215

Realini M.F., González G.E., Font F., Picca P.I., Poggio L., Gottlieb A.M. (2014) Phylogenetic relationships in *Opuntia* (Cactaceae, Opuntioideae) from southern South America. Plant Systematics and Evolution 301: 1123–1134. https://doi.org/10.1007/s00606-014-1154-1

Ritter F. (1979) Kakteen in Südamerica 1. Brazilien/Uruguay/Paraguay. Spangenber, F. Ritter.

Ritter F. (1980) Kakteen in Südamerica 2. Argentinien/Bolivien. Spangenber, F. Ritter.

Salm-Dyck J. (1850) Cacteae in Horto Dyckensi cultae anno 1849. Bonnae. https://doi.org/10.5962/bhl.title.120333

Schumann K. (1890) Cactaceae. In: Martius C.F.P. Flora Brasiliensis Volume 4, Part 2. https://doi.org/10.5962/bhl.title.454

Schumann K. (1899a) Die Cactaceen der Republik Paraguay. Monatsschrift für Kakteenkunde 9: 149–154.

Schumann K. (1899b) Gesamtbeschreibung der Kakteen (Monographia cactacearum). Neudamm, Verlag J. Neumann. https://doi.org/10.5962/bhl.title.10394

SpeciesLink (2019) Digital database, CRIA. Available at http://www.splink.org.br/ [accessed 12 Dec. 2019].

Spegazzini C. (1899) Nova addenda ad floram Patagonigam. Anales de la Sociedad Científica Argentina 48: 44–59.

Spegazzini C. (1901) Contribución al estudio de la flora del Tandil. La Plata, Sesé, Larrañaga y Renovales. https://doi.org/10.5962/bhl.title.9302

Spegazzini C. (1902) Nova addenda ad floram Patagonigam. Anales de la Sociedad Científica Argentina 53: 275–292.

Spegazzini C. (1905) Cactacearum Platensium Tentamen. Anales de la Sociedad Científica Argentina 3(5): 477–521.

Spegazzini C. (1925) Nuevas notas Cactológicas. Anales de la Sociedad Científica Argentina 99: 85–156.

Vega R.R. (2013) Distribución, variación morfológica y correlaciones ecológicas de *Opuntia* Miller (Cactaceae) en Colombia. Montería, Zenú.

